# A translaminar genetic logic for the circuit identity of intracortically-projecting neurons

**DOI:** 10.1101/290395

**Authors:** Esther Klingler, Andres De la Rossa, Sabine Fièvre, Denis Jabaudon

## Abstract

Distinct subtypes of intracortically-projecting neurons (ICPN) are present in all layers, allowing propagation of information within and across cortical columns. How the molecular identities of ICPN relate to their defining anatomical and functional properties is unknown. Here we show that the transcriptional identities of ICPN primarily reflect their input-output connectivities rather than their birth dates or laminar positions. Thus, conserved circuit-related transcriptional programs are at play across cortical layers, which may preserve canonical circuit features across development and evolution.

## Introduction

Neurons of the neocortex are organized into six radial layers, which have appeared at different times during evolution, with the superficial layers representing a more recent acquisition. Input to the neocortex predominantly reaches superficial layers (SL, *i*.*e*. layers (L) 2-4), while output is generated in deep layers (DL, *i*.*e*. L5-6)^1,2^. Intracortical connections, which bridge input and output pathways, are key components of cortical circuits because they allow the propagation and processing of information within the neocortex, thereby transforming afferent signals into cortical output.

Two main types of intracortically-projecting neurons (ICPN) can be distinguished by their axonal features (**Fig. 1a**): (1) excitatory interneurons with short axons projecting locally within cortical columns, which are located in L4 and called spiny stellate neurons (SSN)^3–6^, and (2) excitatory neurons with long axonal projections, including callosally projecting neurons (CPN), which are found in both SL and DL (CPN_SL_ and CPN_DL_)^6–8^. In addition to their distinct axonal features, neurons in these two classes can be distinguished by their hierarchical position within cortical circuits: SSN are the main recipients of thalamic input and project to both CPN and other SSN, while CPN connect with one another, but not back to SSN (**Fig. 1a**)^4,5,9^.

**Figure 1.**
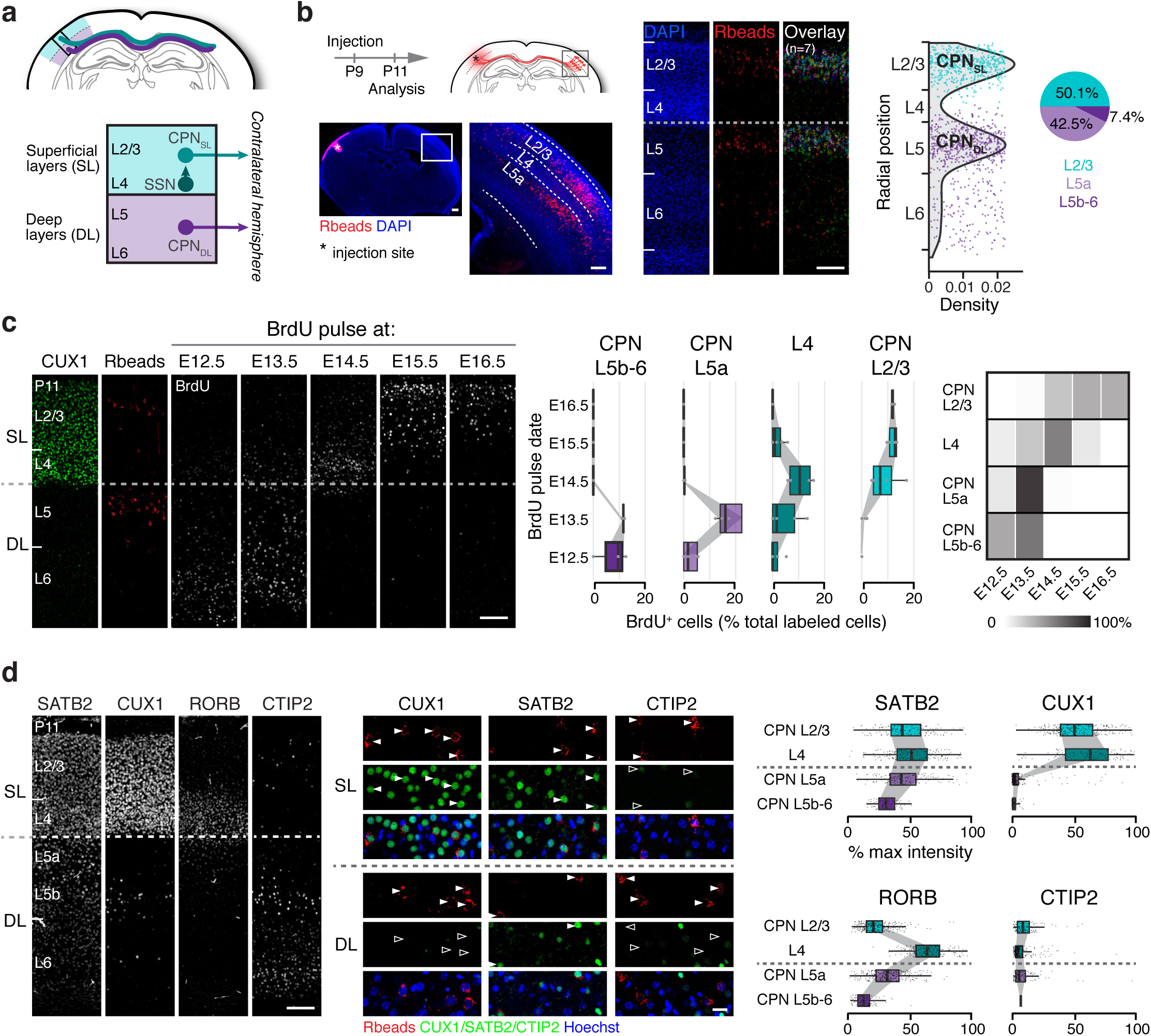
Diversity of intracortically-projecting neurons (ICPN) across cortical laminae. **a**, Schematic representation of the 3 populations of ICPN: callosally-projecting neurons from superficial (CPN_SL_) and deep layers (CPN_DL_) and spiny stellate neurons from layer (L) 4 (SSN). **b**, Distribution of CPN within cortical laminae at P11. CPN were labeled by injection of red Retrobeads (Rbeads) in the contralateral somatosensory cortex. CPN were absent from L4, where SSN are located. **c**, Birthdates of ICPN reflect the inside-out generation of cortical neurons. **d**, Expression of cortical markers by ICPN. Scale bars represent 150 μm (b, c, d, left), 20 μm (d, middle). DL: deep layers, SL: superficial layers.

In this study, we investigate the molecular hallmarks that distinguish SSN, CPN_SL_ and CPN_DL_ and relate their transcriptional signatures with their input-output connectivity (*i*.*e*. their circuit identity). Specifically, taking advantage of the presence of CPN in both SL and DL, we sought to identify lamina-independent genetic hallmarks of a constant circuit motif (*i*.*e*. interhemispheric connectivity) across distinct layers. Using retrograde tracing from the primary somatosensory cortex to label contralateral CPN, we report that CPN_SL_ and CPN_DL_ are born at different times of corticogenesis and have distinct developmental histories. By performing three-way unbiased transcriptomic comparisons between CPN_SL_, CPN_DL_ and SSN, we find that circuit identity supersedes laminar identity in defining ICPN transcriptional diversity. Supporting the functional relevance of a primarily circuit-based transcriptional organization, overexpression of the SSN-specific transcription factor RORB was sufficient to reprogram the circuit identity of CPN within their original layer.

Together, these findings reveal a circuit-based organization of transcriptional programs across cortical layers, which we propose reflects an evolutionary conserved strategy to protect canonical circuit structure (and hence function) across a diverse range of neuroanatomies.

## Results

### ICPN are developmentally and molecularly heterogeneous

In order to characterize the laminar diversity of ICPN, we labeled CPN in the primary somatosensory cortex by injection of fluorescent retrobeads in the contralateral hemisphere (**Fig. 1b**). Retrogradely-labeled cells had a bimodal spatial distribution in SL and DL, and were largely absent from L4, where SSN are located (**Fig. 1b**). This mutually exclusive distribution of SSN and CPN suggests shared lineage relationships between ICPN in which single subtypes are generated at a given time point of corticogenesis.

We next compared the developmental histories of CPN_SL_ and CPN_DL_. Given their distinct laminar location, CPN_DL_ could either be born together with CPN_SL_ and arrest their migration within deep layers, or be born before CPN_SL_, together with other deep-layer neurons. To distinguish between these two possibilities, we determined the birthdates of the distinct ICPN subtypes by performing daily BrdU pulse-injections between embryonic days (E) 12.5 and E16.5, and retrogradely labeled CPN as described above. This approach revealed that CPN_DL_ are born at E12.5 and E13.5 (as are other DL neurons), while CPN_SL_ are mostly born between E15.5 and E16.5 (**Fig. 1c**). Thus, despite similar contralateral projections, CPN_SL_ and CPN_DL_ have non-overlapping developmental (and potentially evolutionary) histories.

We next examined how select markers of distinct types of cortical neurons were expressed across these cells. For example, while SATB2 and CUX1 are strongly expressed by CPN_SL_, but whether CPN_DL_ and SSN also express these genes has not been systematically examined^8,10,11^. Using SATB2, CUX1, RORB (a L4 marker) and CTIP2 (a L5B corticofugal neuron maker) as canonical genes, we report overlap and heterogeneity in gene expression across ICPN (**Fig. 1d**). Most strikingly, while all ICPN expressed SATB2, CPN_DL_ neither expressed CUX1 nor CTIP2 (**Fig. 1d**). Thus, based on this select set of markers, and confirming and extending previous results^8,10^, CPN_DL_, CPN_SL_, and SSN constitute molecularly diverse and partially overlapping populations of cells, which may be linked to their circuit properties.

### Circuit identity supersedes laminar identity in defining ICPN transcriptional diversity

Molecular distinctions between the different types of ICPN could either reflect their laminar identity or their circuit identity (**Fig. 2a**). In the layer-based scenario, CPN_SL_ and CPN_DL_ are only remotely related, reflecting their distinct laminar locations and developmental origins, while in the second scenario, the shared circuit properties of CPN_SL_ and CPN_DL_ are reflected in closely-related transcriptional programs, as has recently been reported in cortical interneurons^12^.

**Figure 2.**
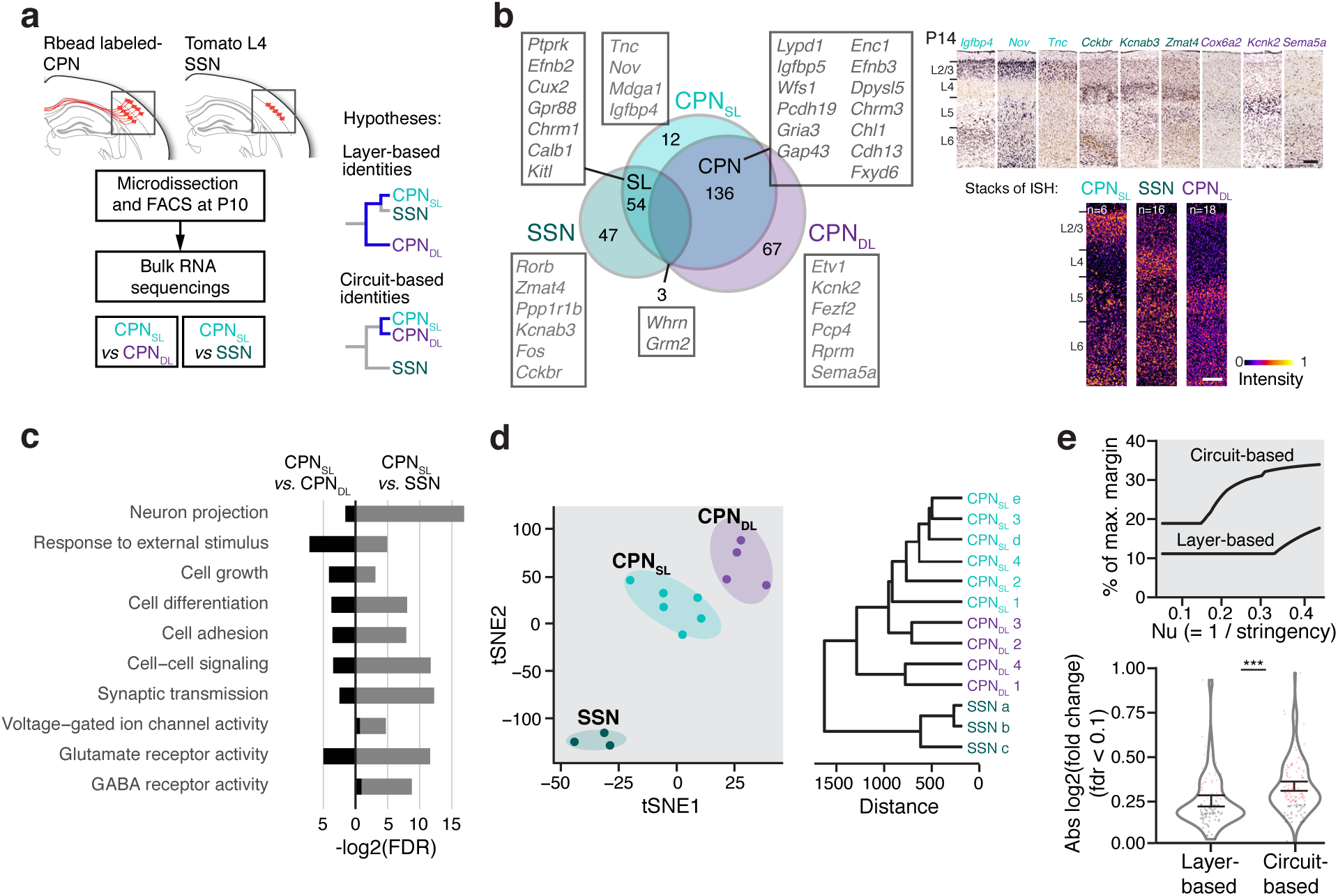
Circuit properties are the primary determinant of transcriptional identity for CPN_SL_, SSN, and CPN_DL_. **a**, Schematic representation of the experimental strategy and working hypotheses. **b**, Three-way transcriptomic comparison between CPN_SL_, SSN, and CPN_DL_ reveals type-specific and shared genes (left). Examples of selected gene *in situ* hybridizations (ISH, source: Allen Brain Atlas) and pseudo-intensities of stacked ISH of all significant genes that were referenced in the database (right). **c**, Ontologies of differentially-expressed genes. **d**, Unbiased clustering delineates CPN_SL_, SSN, and CPN_DL_. Circles represent individual samples. tSNE, t-distributed stochastic neighbour embedding. **e**, SSN *vs*. CPN_SL_ (circuit-based hypothesis) classification is superior to CPN_SL_ *vs*. CPN_DL_ classification (layer-based hypothesis) at all levels of stringency (top). Circuit-specific genes are more differentially expressed than layer-specific genes (bottom). Genes with fold changes > 2 are highlighted in red. \*\*\**P*<0.001; Welch’s two-sample t-test. Scale bar in **b** represents 150 μm.

To distinguish between these two possibilities, we examined the genetic signatures of CPN_SL_, CPN_DL_, and SSN using RNA sequencing (**Fig. 2a**). First, we isolated CPN_SL_ and CPN_DL_ by using retrograde-labeling, laminar microdissection, and fluorescence activated cell sorting (FACS) at P10, a time at which interhemispheric connectivity is largely achieved^13^. Second, we isolated CPN_SL_ from SSN by performing retrograde labeling and FACS in transgenic mice in which SSN specifically express tdTomato (*Scnn1aTg3*^Cre^*xAi14tdT* mice^5^). Two-way transcriptional analyses comparing CPN_SL_ and CPN_DL_ identified 136 differentially-expressed genes, while comparison of CPN_SL_ and SSN identified 204 differentially-expressed genes (**Fig. 2b** left, and **Supplementary Fig. 1** and **2 and Supplementary Table 1**), whose specificities were confirmed with *in situ* hybridization data (**Fig. 2b** right, and **Supplementary Fig. 3**). Identified genes included previously known markers such as *Mdga1* and *Pcp4*, which were enriched in CPN_SL_ and CPN_DL_ respectively, and *Rorb*, which was enriched in SSN (**Supplementary Fig. 3**). Interestingly, corticofugal neuron markers such as *Fezf2* and *Bcl11b* were weakly expressed in CPN_DL_, consistent with the presence of striatal projections in at least a subset of these neurons^14,15^. Supporting the functional relevance of these transcripts, comparison of gene ontologies identified greater enrichment in transcripts related to neuron projections and activity/physiology-related functions when comparing CPN_SL_ with SSN, consistent with the distinctive circuit position and function of SSN within cortical circuits (**Fig. 2c)**.

To assess the transcriptional relationship between CPN_SL_, CPN_DL_, and SSL, we performed unsupervised clustering of samples based on transcriptional signatures. Hierarchical clustering revealed that CPN_SL_ and CPN_DL_ are more closely related to one another than to SSN (**Fig. 2d**). This suggests a primarily circuit-based organization of transcriptional programs. To formally demonstrate this possibility, we compared the discriminative power of the layer-based taxonomy to a circuit-based taxonomy, as previously described^16^. This quantitative assessment of these two taxonomies revealed that the circuit-based classification was more discriminative than the layer-based classification at all levels of stringencies examined (**Fig. 2e**, top). Accordingly, cell-type specific genes were more differentially expressed between CPN_SL_ and SSN than between CPN_SL_ and CPN_DL_ (**Fig. 2e**, bottom). Together, these data indicate that ICPN molecular identities more closely correspond to their circuit properties than laminar location.

### RORB-overexpressing CPN_SL_ acquire SSN-like circuit properties

Finally, we sought to identify a functional molecular counterpart to the circuit-based classification identified above, using *Rorb* as a proof-of-principle transcript. Indeed, this orphan receptor shows a 3-fold enrichment in SSN *vs*. CPN_SL_ (**Supplementary Table 1**) and has been implicated in circuit assembly within and beyond the cortex^17–19^. Here, we directly examined the function of RORB in intracortical circuit assembly by assessing whether targeted overexpression in ICPN induces acquisition of SSN-type morphology, electrophysiology, and circuit connectivity. For this purpose, we electroporated a plasmid coding for RORB at E16.5, the time of birth of CPN_SL_. As previously reported, a fraction of RORB-overexpressing cells showed migratory defects and did not reach the cortex^17^ (**Supplementary Fig. 4**), yet some cells maintained their normal migration and reached superficial layers (ICPN_RORB_, **Fig. 3a**, left and **Supplementary Fig. 4**). We first compared the morphology of ICPN_RORB_ with control cells born at E14.5 (SSN) or at E16.5 (ICPN_L2/3_). In contrast to ICPN_L2/3_, the SSN are characterized by the absence of an apical dendrite^3,5,20^. Strikingly, as it is the case for SSN, ICPN_RORB_ actively retracted their apical dendrite between P3 and P7, which was not the case in control ICPN_L2/3_ (**Fig. 3a**, center and right and **Supplementary Fig. 4**). Thus, RORB expression controls acquisition of a key morphological feature of SSN.

**Figure 3.**
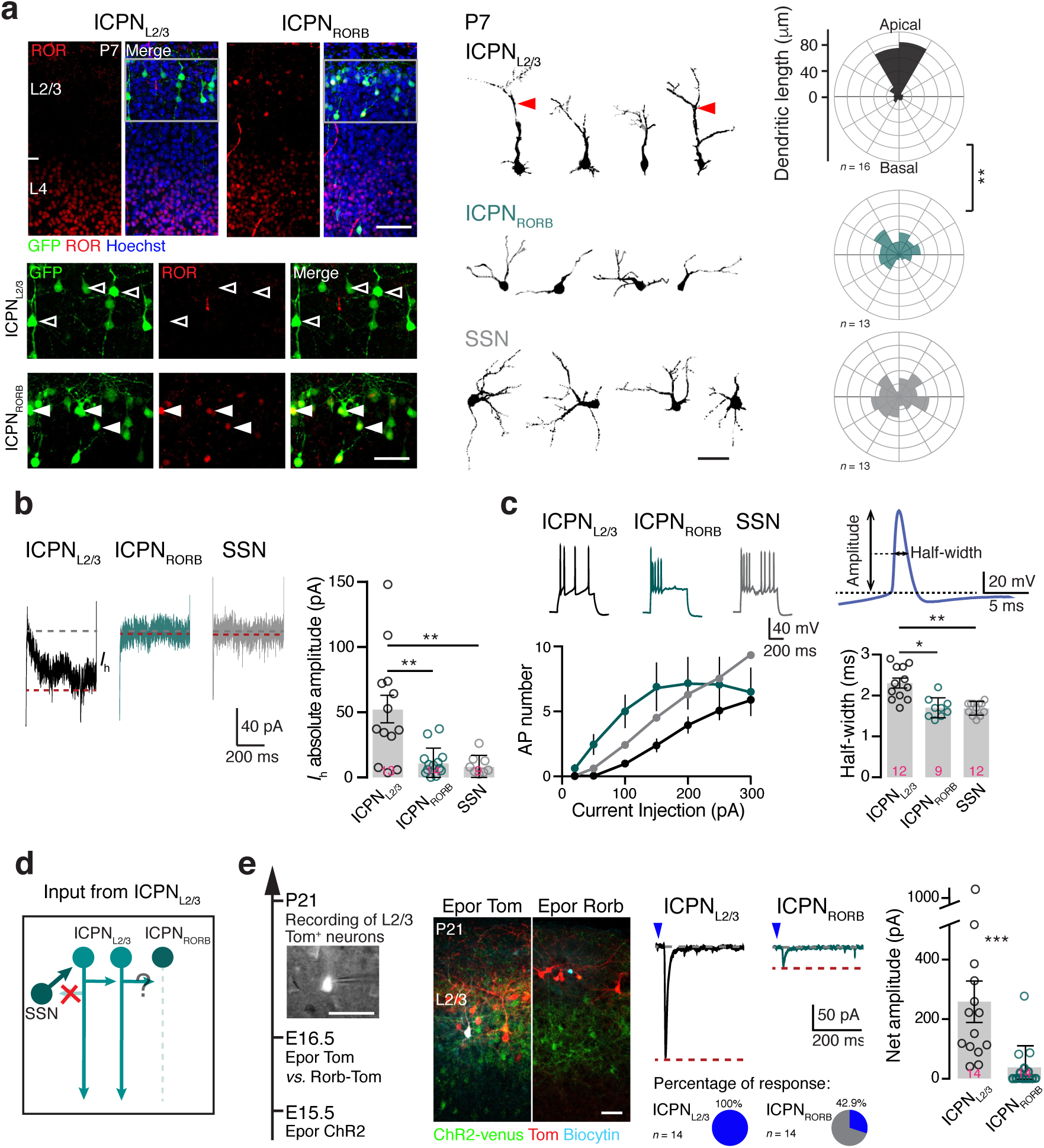
ICPN_RORB_ acquire SSN-like morphology and electrophysiological/circuit properties. **a**, In contrast to ICPN_L2/3_, RORB-overexpressing ICPN_L2/3_ (ICPN_RORB_, filled arrowheads) and SSN do not have an apical dendrite at P7. **b**, In contrast to ICPN_L2/3_, ICPN_RORB_ and SSN lack *I*_*h*_ currents. Sample traces show *I*_h_ current in response to a 500 ms, -40 mV square voltage step. **c**, ICPN_RORB_ acquire SSN-like excitability. **d**, Schematic representation of L2/3-L4 canonical microcircuit. **e**, ICPN_RORB_ display SSN-like cortical circuit features. Left, experimental design and illustration. Right, sample traces, percentage of response and response amplitude following 0.5 ms light stimulation of ICPN_L2/3_ (blue arrowhead). Scale bars represent 50 μm (a, top, f), 30 μm (a, bottom), 25 μm (b).

We next examined whether ICPN_RORB_ also acquired electrophysiological properties of SSN, including changes in *I*_h_-type cationic conductances and membrane excitability^5,21^. Consistent with acquisition of SSN-type electrophysiological features, whole-cell patch-clamp recordings in acute cortical slices showed a lack of *I*_h_ currents in ICPN_RORB_, as well as increased membrane excitability (**Fig. 3b**,**c**). Interestingly, action potential duration was shorter in both SSN and ICPN_RORB_ compared to ICPN_L2/3_, which could account for the higher firing rate in the former cells (**Fig. 3c**). Thus, RORB expression controls acquisition of key electrophysiological features of SSN.

Finally, we examined whether ICPN_RORB_ acquired an SSN-type circuit identity. Consistent with acquisition of SSN-type local axonal projections, long-range projections were lacking in ICPN_RORB_, as previously reported (**Supplementary Fig. 4**)^18^. Focusing on local microcircuit properties, we next examined the local connectivity of ICPN_RORB_. ICPN_L2/3_ normally receive strong input from other ICPN_L2/3_, while SSN_L4_ do not receive ICPN_L2/3_ input^4,5,9^. We therefore examined whether ICPN_RORB_ displayed SSN-type input properties, *i*.*e*. lacked ICPN_L2/3_ input (**Fig. 3d**). To this end, we targeted channelrhodopsin 2 (ChR2) expression into deep ICPN_L2/3_ *via in utero* electroporation at E15.5 and recorded photo-induced post-synaptic responses in superficial E16.5-born ICPN_L2/3_. In contrast to ChR2^−^ ICPN_L2/3_ neurons, which all displayed synaptic responses following optogenetic stimulation of homotypic neurons, only 6/14 ICPN_RORB_ responded to ICPN_L2/3_ stimulation, with dramatically smaller amplitudes than in control cells (**Fig. 3e**). Together, these findings reveal acquisition of SSN-type morphological, electrophysiological and circuit properties by ICPN_RORB_.

Our findings reveal a genetic organization of ICPN in which transcriptional programs more closely reflect circuit properties than laminar location or developmental origins. From a phylogenetic perspective, we find these results interesting considering that superficial cortical layers are a recent evolutionary acquisition of mammals^1,2^: this suggests either that a specialized progenitor class generates CPN throughout corticogenesis, or that a convergent evolution has occurred in deep and superficial layer neurons, in which similar molecular programs were selected for trans-callosal axon extension. Finally, using RORB as a proof-of-principle transcript, we show that ectopic expression of a single gene is sufficient to orchestrate the coordinated acquisition of the morphological, physiological and circuit properties of another intracortically-projecting neuron subtype. Along with similar recent findings in inhibitory interneurons^12^, this suggests that circuit properties are critical end-point determinants of neuronal identity and the result of convergent molecular programs during neuronal differentiation.

## Acknowledgements

We thank the Genomics Platform and FACS Facility of the University of Geneva, A. Benoit and T. Kastylevsky for technical assistance; and members of the Jabaudon laboratory as well as U. Tomasello, G. Limoni, I. Vitali and E. Azim for constructive comments on the manuscript. The Jabaudon Laboratory is supported by the Swiss National Science Foundation and the Brain and Behavior foundation; E.K. is supported by a grant from the Machaon Foundation.

## Author contributions

E.K., A.D.l.R., and D.J. conceived the project and designed the experiments. E.K., A.D.l.R and S.F. performed the experiments. E.K. and D.J. wrote the manuscript.

## Materials and methods

### Mouse strains

C57Bl/6 male and female pups and adult mice were used. The *Scnn1a-cre* mouse line (Jackson Laboratories^22^; #009613) was crossed with CAG-tdTomato reporter mice (Jackson Laboratories; #007914). All experimental procedures were approved by the Geneva Cantonal Veterinary Authority and conducted according to the Swiss guidelines.

### Plasmids

We generated plasmids using a standard endotoxin-free Qiagen kit (#12362). The ChR2 T159C^23^ plasmid was subcloned into the pCAGIG_IRES_GFP vector. The pCBIG_Rorb_IRES_GFP plasmid was obtained from Addgene (#48709)^17^.

### *In utero* electroporation

Timed pregnant C57Bl6/J mice (Charles River Laboratory) were electroporated *in utero* at E14.5, E15.5 or E16.5 as previously described^17^.

### BrdU pulse labeling

A single dose of 50 mg/kg of animal weight of BrdU (16 mg/ml) was administered by an intraperitoneal injection in the mother during pregnancy, from embryonic day (E) 12.5 to E16.5.

### Retrograde labeling

Anesthetized pups were placed in a stereotaxic apparatus at postnatal day (P) 9 and injected with red Retrobeads™ IX from Lumafluor (for CPN_SL_ *vs*. CPN_DL_ comparison) or with Alexa-488 conjugated cholera toxin subunit B (CTB, Invitrogen, #C-34775) (for CPN_SL_ *vs*. SSN comparison) in S1 (200 nl; coordinates from the lambda: anteroposterior: 3 mm, mediolateral: 3mm).

### Immunohistochemistry

Postnatal mice were perfused with 4% paraformaldehyde (PFA) and brains were fixed overnight in 4% PFA at 4 °C. Eighty μm vibrating microtome-cut coronal sections (Leica, VT1000S) were incubated 2h at room temperature in a blocking/permeabilizing solution containing 5% bovine serum albumin and 0.3% triton X-100 in PBS, and incubated for 2 days with primary antibodies at 4°C. Sections were then rinsed three times in PBS and incubated with the corresponding Alexa-conjugated secondary antibodies (1:500; Invitrogen) for 2 h at room temperature. For immunohistochemistry against RORB, a pre-treatment in 10 mM sodium citrate, 0.05% tween 20 buffer (pH6) at 85°C for 40 minutes was performed before running the same protocol mentioned above of the procedure. For immunohistochemistry against BrdU, a pre-treatment with 2N HCl at 37°C for 40 minutes was performed. Primary antibodies and their dilutions were: mouse anti-human ROR (Perseus proteomics, # PP-H3925-00, 1:200), chicken anti-GFP (Invitrogen, # A10262, 1:2000), rat anti-BrdU (Abcam, # ab6326, 1:200), rat anti-CTIP2 (Abcam, # ab18465, 1:500), rabbit anti-CUX1 (Santa Cruz, # sc13024, 1:250), mouse anti-SATB2 (Abcam, # ab51502, 1:200).

### Image acquisition and quantifications

All images were acquired on a Nikon A1r spectral confocal microscope, equipped with 40x 0.6 CFI ELWD S Plan Fluor WD objective. Cells were selected for analysis only if there was no overlap with the primary dendrites of other labeled neurons. GFP/RORB overexpressing neurons were reconstructed for quantitative analysis of neuronal dendritic arborization using Imaris software. Quantifications of CUX1/SATB2/RORB/CTIP2 fluorescence intensity were done using Fiji software, by normalizing the row values to the background intensity (measured on a region of the section without positive cell) on each section and displayed them as the percent of max value on the section. For CTIP2 intensity quantification, the max value was measured in a positive L5 cell (since most ICPN express very low to not detectable levels of CTIP2).

To quantify BrdU^**+**^ CPN after injection of BrdU at different embryonic stages, we calculated the percentage of Retrobead-labeled CPN in L2/3, L5a, and L5-6 that displayed a high intensity of BrdU. Only sections with at least 10 Retrobead-labeled CPN per layer were kept for analyses (n=2 to 3 sections per pup, n=3 to 5 pups per age). For L4 BrdU^**+**^ cells, we delineated L4 with CUX1 staining, from which the barrels were identifiable, and quantified the percentage of BrdU^**+**^ cells in this area. For the heatmap in Fig 1c, the total number of BrdU^**+**^ ICPN was normalized to 100% per layer, showing the peak of birth for each ICPN population per layer. Error bars represent s.e.m.

### Tissue microdissection, cell sorting and RNA sequencing

Microdissection of primary somatosensory cortex (S1) from one litter (4 pups) corresponds to one biological replicate (*n* = 3 to 6 for each condition). Fresh coronal brain sections (600 μm) were cut on a vibrating microtome (Leica, VT1000S) and S1 was microdissected using a Leica Dissecting Microscope (Leica, M165FC) in ice-cold oxygenated artificial cerebrospinal fluid (ACSF) under RNase-free conditions. For the comparison of SSN *vs*. CTB-labeled CPN_SL_, we used the same procedure as in ref. 10 for cell dissociation. For the comparison of Retrobead-labeled CPN_SL_ *vs*. CPN_DL_, cells were dissociated by incubating micro-dissected samples in 0.5 mg/mL pronase (Sigma, #P5147) at 37°C for 10 minutes, followed by incubation in 5% bovine serum albumin for 3 minutes and manual trituration in ACSF using pulled glass pipettes. Cells were then centrifuged for 10 minutes at 600 rpm and resuspended before filtration using a 70 μm cell strainer (ClearLine, # 141379C). Cells were then incubated for 10 minutes at 37°C with Hoechst (0.1 mg/mL) and isolated using a Beckman Coulter Moflo Astrios FAC-sorter. Singlet Hoechst^+^ cells were sorted according to their Forward and Side scattering properties (FSC and SSC), and their negativity for Draq7TM (Viability dye, Far red DNA intercalating agent, Beckman Coulter, #B25595). RNA was extracted using an RNeasy kit (Qiagen, #74034). cDNA libraries were obtained using SMARTseq v4 kit (Clontech, # 634888) and sequenced using HiSeq 2500 sequencer.

### Analyses

For bulk RNA sequencing data analyses, we kept only genes expressed more than 20 rpm in at least one of sample. To perform three-way analyses on the layer *vs*. circuit datasets, we normalized the expression of genes by the means of CPN_SL_ samples from both experiments. For Fig.2d, samples with similar expression of genes and therefore similar principal components loadings, are most likely to localize near each other in the embedding^24,25^. Hierarchical clustering was performed using euclidian distance metrics. To compare the discriminative power of the layer-based classification (that is, SL *vs*. DL) with the circuit-based classification (that is, local *vs*. callosal), we used the same approach as described in ref. 16 (the paragraph below is directly modified from the original description in this study): we trained 2 linear nu-support vector machine (nu-SVM) classification models. Nu corresponds to the degrees of freedom of the SVM model, and thus inversely correlates with stringency. We determined the maximal margin of separation between the two populations (that is, SL *vs*. DL CPN, or SSN *vs*. CPN_SL_), which indicates how distinct these two populations are. Because the ‘nu’ parameter controls the stringency of the model, we confirmed the results using a range of nu values between 0.1 and 0.5. We looked for genes differentially expressed in the layer and the circuit models using SVM with nu=0.3. We considered as differentially expressed genes with a FDR < 0.1.

Statistical analyses of morphological and electrophysiological parameters were performed using Graphpad Prism software. For statistical analyses of neuron morphology at P7, we used 2-way ANOVA and multiple comparisons with Bonferroni correction (n=17 ICPN_L2/3_; n=13 ICPN_RORB_; n=13 SSN_L4_). For analyses of electrophysiological parameters, we used Kruskal-Wallis non-parametric test with multiple comparisons when comparing the 3 ICPN populations and Mann-Whitney test when comparing only ICPN_L2/3_ and ICPN_RORB_. The number of recorded cells is indicated on each figure. For statistical analyses of dendritic length at P21, we confirmed normality of the data using D’Agostino & Pearson normality test and performed unpaired t-test.

### Electrophysiology

300 µm thick coronal slices from P21 mice were cut in cooled ACSF containing 125 mM NaCl, 2.5 mM KCl, 1 mM MgCl, 2.5 mM CaCl_2_, 1.25 mM Na_2_HPO_4_, 26 mM NaHCO_3_ and 11 mM glucose, oxygenated with 95% O_2_ and 5% CO_2_. Slices were kept at room temperature and allowed to recover for 1h before recording. Under low magnification, the barrels in L4 could be readily identified, and high-power magnification was used to guide the recording electrode onto visually identified neurons. Whole-cell voltage-clamp recordings were performed with 3–4 MΩ electrodes filled with a solution containing 140 mM KCH_3_SO_3_, 2 mM MgCl_2_, 4 mM NaCl, 5 mM P-creatine, 3 mM Na_2_ATP, 0.33 mM GTP, 0.2 mM EGTA and 10 mM HEPES adjust to 300 mOsm l-1 1 and pH 7.2 with KOH. Currents were amplified (Multiclamp 700B, Axon Instruments), filtered at 5 kHz and digitized at 20 kHz (National Instruments Board PCI-MIO-16E4, Igor, WaveMetrics). *I*_h_ was measured in voltage clamp using a -40mV step (500 ms) and was calculated by the difference of current between the beginning and the end of the voltage step. The firing pattern was studied by current clamp recording of the neurons during the injection of 20 to 300 pA of current (50 pA stepped) for 500 ms. Optogenetic stimulations were performed using 0.5 ms blue LED light pulses at 0.05 Hz in L2/3 ChR2-expressing sections, and photo-induced EPSCs were recorded from ICPN_L2/3_ and ICPN_RORB_ in presence of 10 µM bicuculline. No series resistance compensation was used. Values are presented as mean ± s.e.m.

---

**Supplementary Fig. 1.**
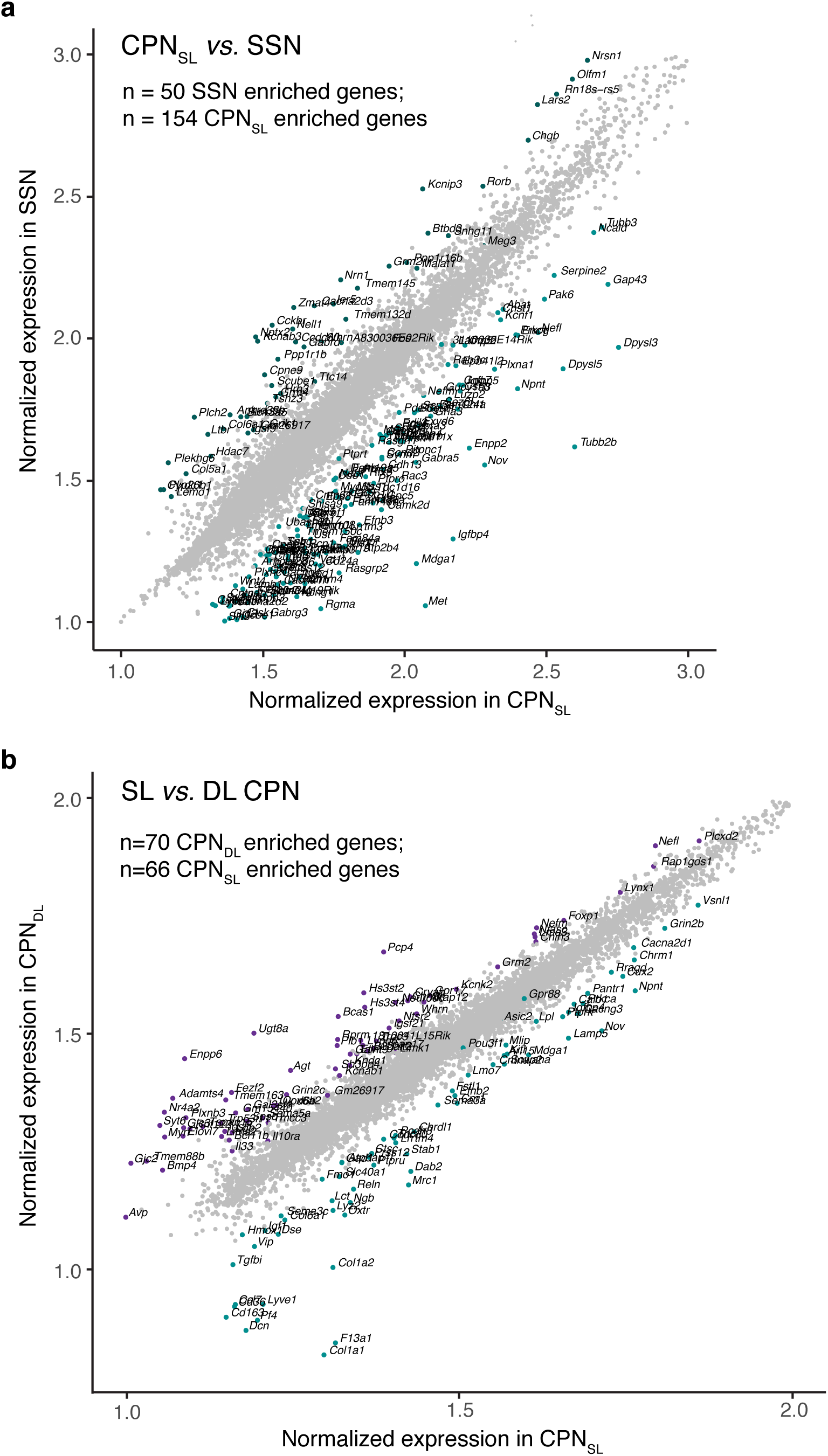
Differential gene expression in CPNSL *vs*. SSN (a), and in SL *vs*. DL CPN (b). Annotated genes are significantly differentially expressed between the populations.

**Supplementary Fig. 2.**
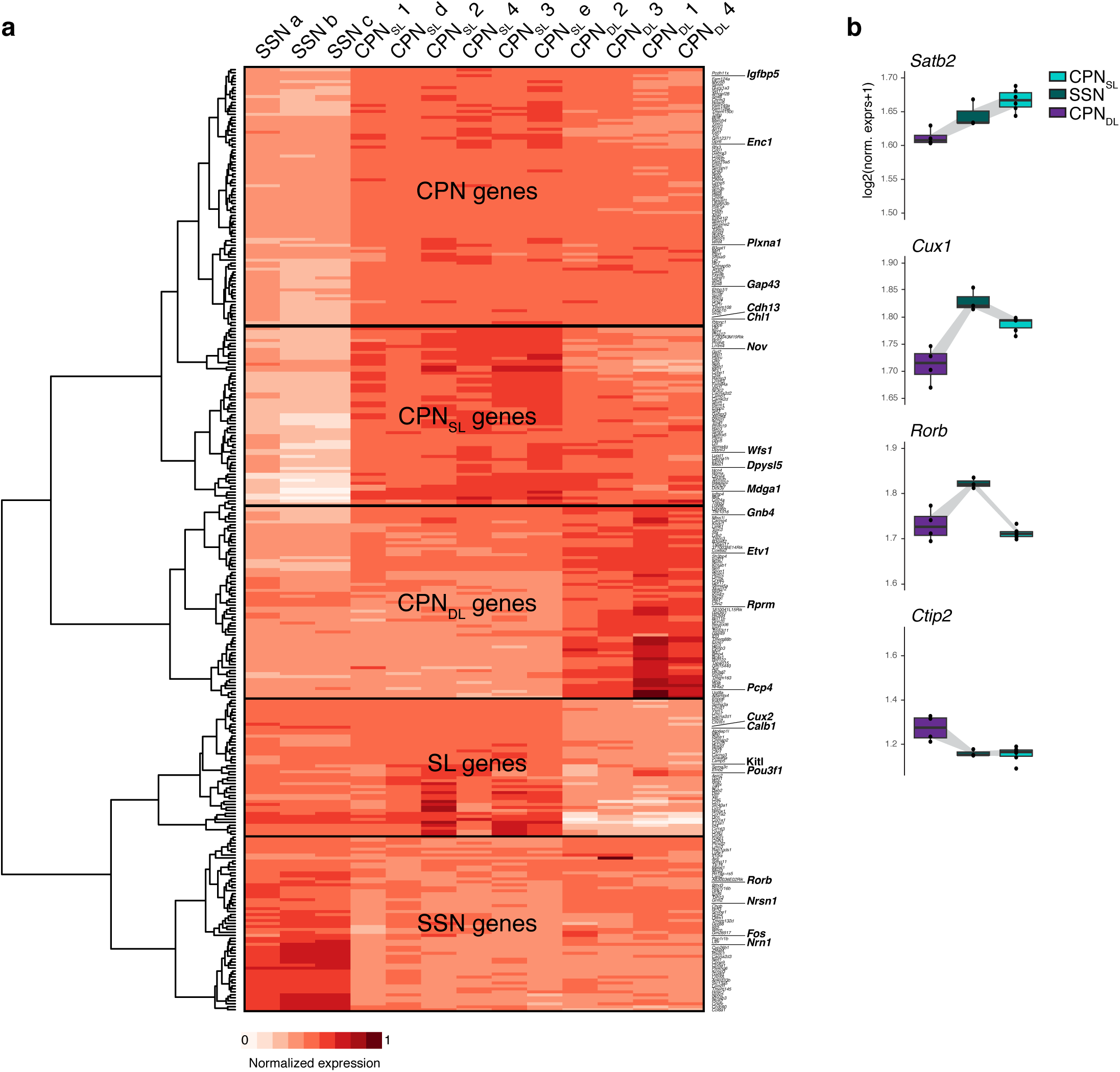
ICPN differentially expressed genes. **a**, Expression heatmap and unbiased clustering of all differentially expressed genes in CPN_SL_ *vs*. SSN and SL *vs*. DL CPN datasets. **b**, Expression of the markers used for primary characterization of the 3 ICPN populations (compare with values for corresponding protein expression in Fig. 1d).

**Supplementary Fig. 3.**
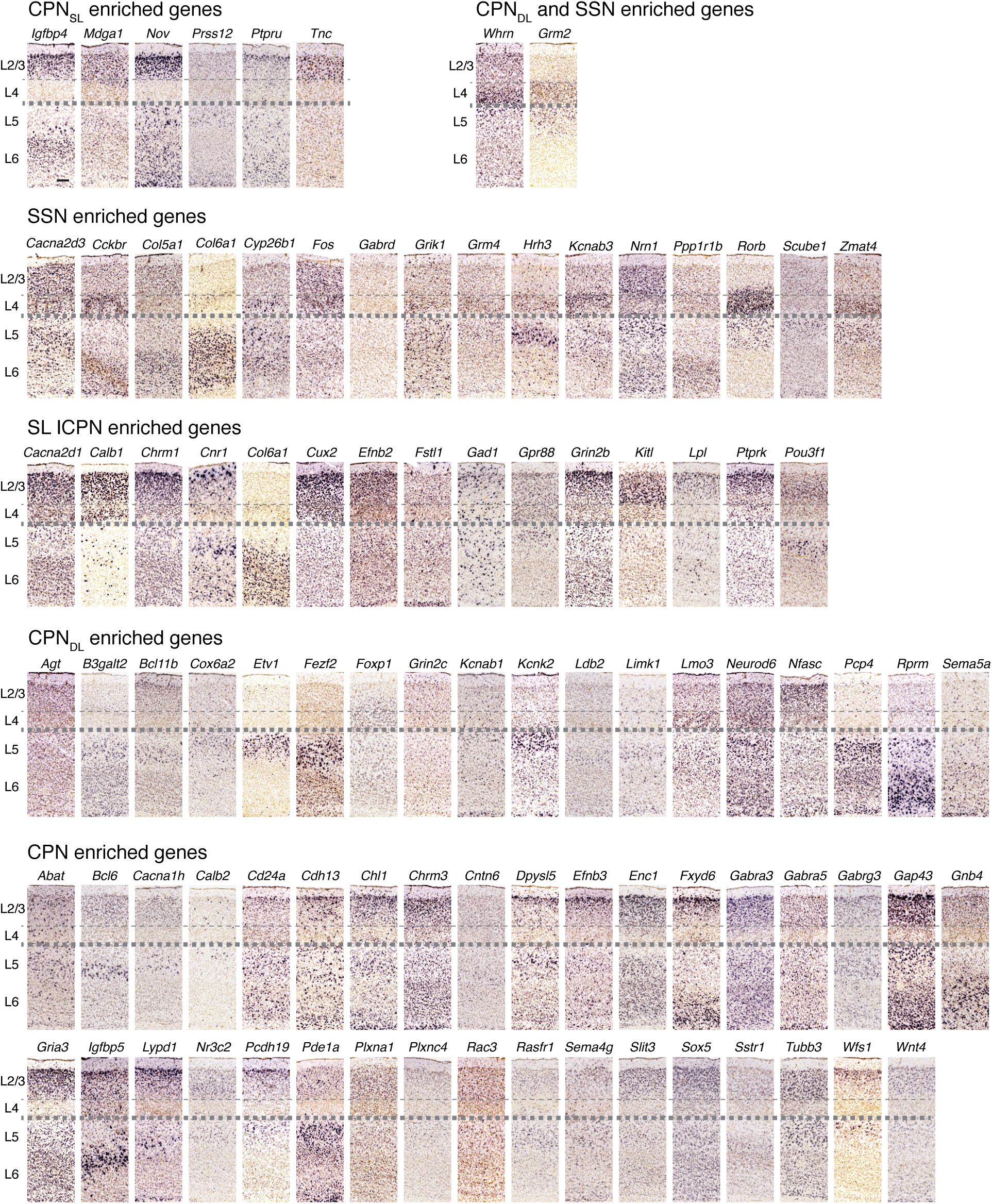
Expression of differentially expressed genes by ICPN. Data from Allen Brain Atlas database (http://developingmouse.brain-map.org/). Scale bar represents 100 μm.

**Supplementary Fig. 4.**
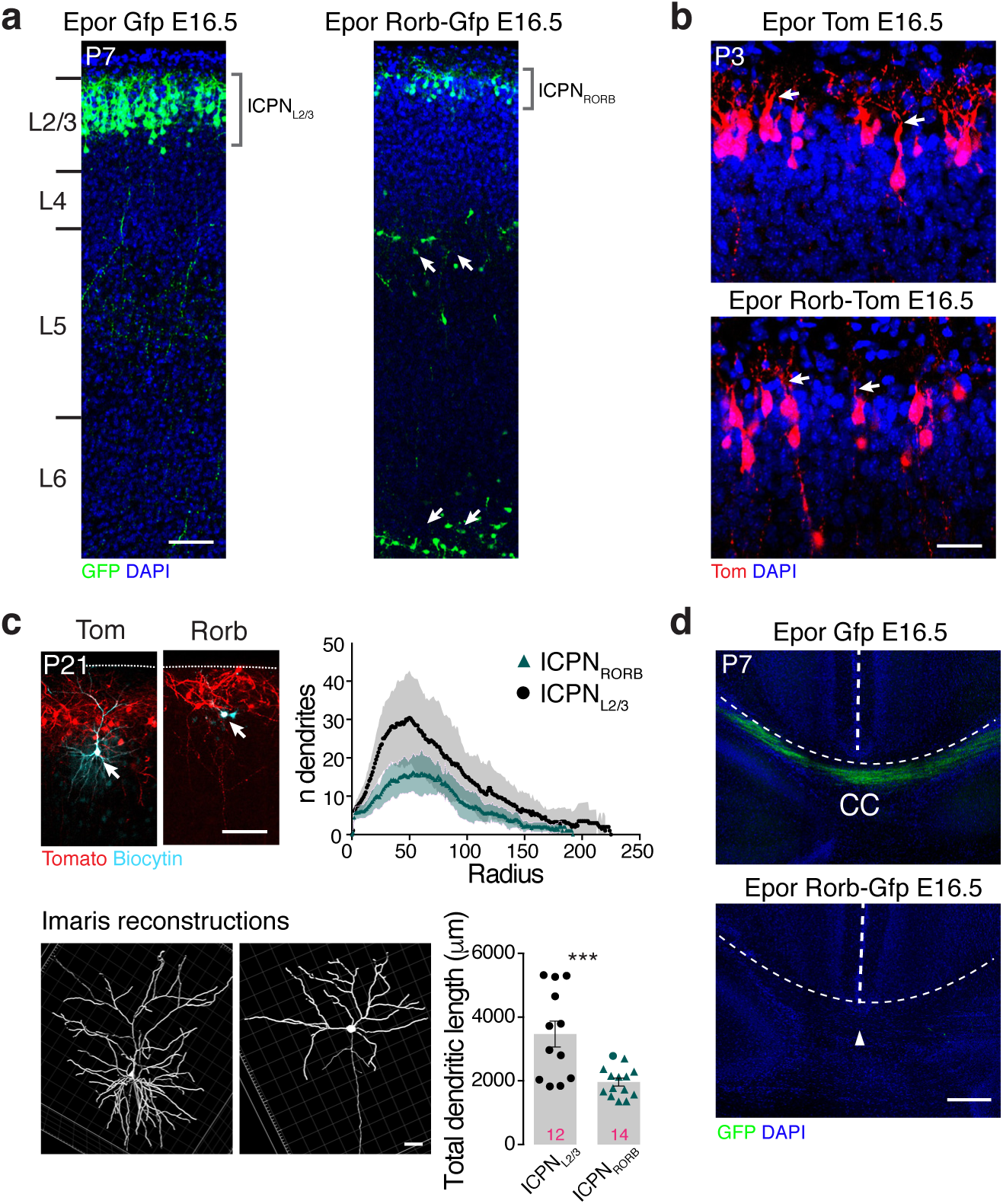
ICPN_RORB_ aquire SSN-like morphology and circuit properties. **a**, ICPN_L2/3_ labeled by GFP after *in utero* electroporation at E16.5. A fraction of RORB overexpressing cells are abnormally positioned (arrows), though a significant proportion reaches L2/3 (ICPN_RORB_). **b**, At P3, ICPN_RORB_ display an apical dendrite (arrows) as do ICPN_L2/3_. **c**, In adults, ICPN_RORB_ lack an apical dendrite and have a reduced number of dendrites as well as a reduced dendritic length compared to ICPN_L2/3_. Imaris reconstructions were performed on biocytin filled electroporated cells, followed by Sholl analyses. **d**, At P7, ICPN_RORB_ do not extend an axon through the corpus callosum (CC, arrowhead) in contrast to control ICPN_L2/3_. Scale bars represent 150 μm (**a, b, d**, top), 40 μm (**c**), 20 μm (**d**, bottom).

